# The lawful imprecision of human surface tilt estimation in natural scenes

**DOI:** 10.1101/180984

**Authors:** Seha Kim, Johannes Burge

**Affiliations:** Department of Psychology, University of Pennsylvania; Neuroscience Graduate Group, University of Pennsylvania; Bioengineering Graduate Group, University of Pennsylvania

## Abstract

Estimating local surface orientation (slant and tilt) is fundamental to recovering the three-dimensional structure of the environment, but it is unknown how well humans perform this task in natural scenes. Here, with a high-fidelity database of natural stereo-images with groundtruth surface orientation at each pixel, we find dramatic differences in human tilt estimation with natural and artificial stimuli. With artificial stimuli, estimates are precise and unbiased. With natural stimuli, estimates are imprecise and strongly biased. An image-computable normative model grounded in natural scene statistics predicts human bias, precision, and trial-by-trial errors without fitting parameters to the human data. These similarities suggest that the complex human performance patterns with natural stimuli are lawful, and that human visual systems have internalized local image and scene statistics to optimally infer the three-dimensional structure of the environment. The current results help generalize our understanding of human vision from the lab to the real world.

## Introduction

Understanding how vision works in natural conditions is a primary goal of vision research. One measure of success is the degree to which performance in a fundamental visual task can be predicted directly from image data. Estimating the 3D structure of the environment from 2D retinal images is just such a task. However, relatively little is known about how the human visual system estimates 3D surface orientation in natural scenes.

3D surface orientation is typically parameterized by slant and tilt. Slant is the amount by which a surface is rotated away from an observer; tilt is the direction of rotation in which the surface is slanted (Fig. 1A). Compared to slant, tilt has received little attention, even though both are critically important for successful interaction with the 3D environment. Even if slant has been accurately estimated, humans must estimate tilt to determine where they can walk. Surface with tilts of 90°, like the ground plane, can sometimes be walked on. Surfaces with tilts of 0° or 180°, like the sides of tree trunks, can never be walked on.

**Figure 1.**
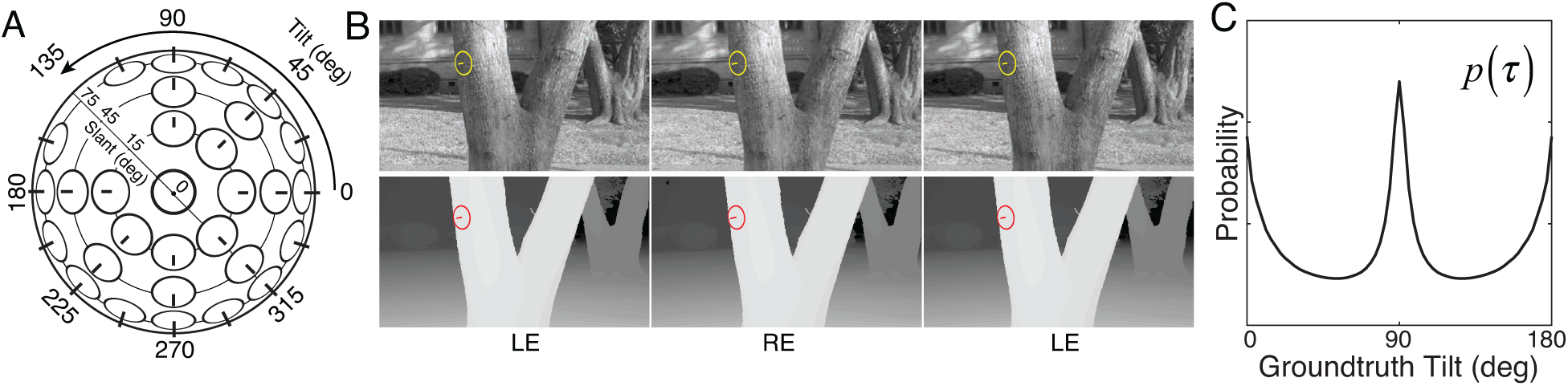
Tilt and slant, natural scene database, and tilt prior. **A** Tilt is the direction of slant. Slant is amount of rotation out of the reference (e.g. frontoparallel) plane. **B** Example stereo-image pair (top) and corresponding stereo-range data (bottom). The gauge figure indicates local surface orientation. To see the scene in stereo 3D, free-fuse the left-eye (LE) and right-eye (RE) images. **C** Prior distribution of unsigned tilt in natural scenes, computed from 600 million groundtruth tilt samples in the natural scene database (see Methods). Cardinal surface tilts associated with the ground plane (90°) and tree trunks (0° and 180°) occur far more frequently than oblique tilts in natural scenes. Unsigned tilt, *τ* = [0°, 180°), indicates 3D surface orientation up to a sign ambiguity (i.e. tilt modulo 180°).

Numerous psychophysical, computational, and neurophysiological studies have probed the human ability to estimate surface slant, surface tilt, and 3D shape [1–20]. Lawful performance has been observed and models have been developed that nicely describe performance. However, the surface shapes (e.g. planar) and artificial surface markings (e.g. 1/f texture) used in most studies are not particularly representative of the variety of surface shapes and markings encountered in natural viewing; surfaces in natural scenes are often curved or rough and are marked by more complicated surface textures. Thus, performance with simple artificial scenes may not be representative of performance with natural scenes. Also, models developed with artificial scenes often generalize poorly (or cannot even be applied) to natural scenes. These issues concern not just studies of 3D surface orientation perception but vision and visual neuroscience at large.

Few studies have examined the human ability to estimate 3D surface orientation with natural photographic images, the stimuli that our visual systems evolved to process. None, to our knowledge, have done so with high-resolution groundtruth surface orientation information. There are good reasons for this gap in the literature. Natural images are complex and difficult to characterize mathematically, and groundtruth data about natural scenes is notoriously difficult to collect. Research with natural stimuli has often been criticized (justifiably) on the grounds that natural stimuli are too complicated or too poorly controlled for strong conclusions to be drawn from the results. The challenge, then, is to develop experimental methods and computational models that can be used with natural stimuli without sacrificing rigor and interpretability.

Here, we report an extensive examination of human 3D tilt estimation from local image information with natural stimuli. We sampled thousands of natural image patches from a recently collected stereo-image database of natural scenes with precisely co-registered distance data (Fig. 1B) [21]. Groundtruth surface orientation was computed directly from the distance data (see Methods). Human observers binocularly viewed the natural patches and estimated the tilt at the center of each patch. The same human observers also viewed conventional artificially textured planar stimuli matched to the groundtruth tilt, slant, distance, and luminance contrast of the natural stimuli. Then, we compared human performance to the predictions of a normative image-computable model that makes the best possible use of the available image information for the task. This experimental design enables direct, meaningful comparison of human performance across stimulus types, allowing one to isolate important stimulus differences and to interpret human response patterns with respect to principled predictions provided by the model.

A rich set of results emerges. First, tilt estimation in natural scenes is hard; compared to performance with artificial stimuli, performance with natural stimuli is poor. Second, with natural stimuli, human tilt estimates cluster at the cardinal tilts (0°, 90°, 180° and 270°), echoing the prior distribution of tilts in natural scenes (Fig. 1C) [21–23]. Third, human estimates tend to be more biased and variable when the groundtruth tilts are oblique (e.g. 45°). Fourth, at each groundtruth tilt, the distributions of human and model errors tend to be very similar, even though the error distributions themselves are highly irregular. Fifth, human and model observer trial-by-trial errors are correlated, suggesting that similar (or strongly correlated) stimulus properties drive both human and ideal performance. Sixth, differences in estimation performance with natural and artificial stimuli can be largely attributed to local depth variation, a pervasive performance-altering feature of natural scenes that is not explicitly considered in most investigations. Together, these results represent an important step towards the goal of being able to predict human percepts of 3D structure directly from photographic images in a fundamental natural task.

## Results

Human observers binocularly viewed thousands of randomly sampled patches of natural scene; they viewed an equal number of stimuli at each of 24 tilt bins between 0 and 360°. The stimuli were presented on a large (2.0x1.2m) stereo front-projection system 3m from the observer. This relatively long viewing distance minimizes focus cues to flatness. Except for focus cues, the display system recreates the retinal images that would have been formed by the original scene. Each scene was viewed binocularly through a small virtual aperture (1° or 3°) positioned 5 arcmin of disparity in front of the sampled point in the scene (Fig. 2A); the viewing situation is akin to looking at the world through a straw [24]. Observers reported, via a mouse-controlled probe, the estimated surface tilt at the center of each patch (Fig. 2B). We pooled data across human observers and aperture sizes: with unsigned tilt (signed tilt modulo 180°), tilt estimation performance was similar for all observers and aperture sizes (Fig. S1, S2). The same observers also estimated surface tilt with an extensive set of artificial planar stimuli that were matched to the tilts, slants, distances, and contrasts of the natural stimuli presented in the experiment. Thus, any observed performance differences between natural and artificial stimuli cannot be attributed to these dimensions. The impact of slant and distance on tilt estimation is similar with both stimulus types (Fig. 2C).

**Figure 2.**
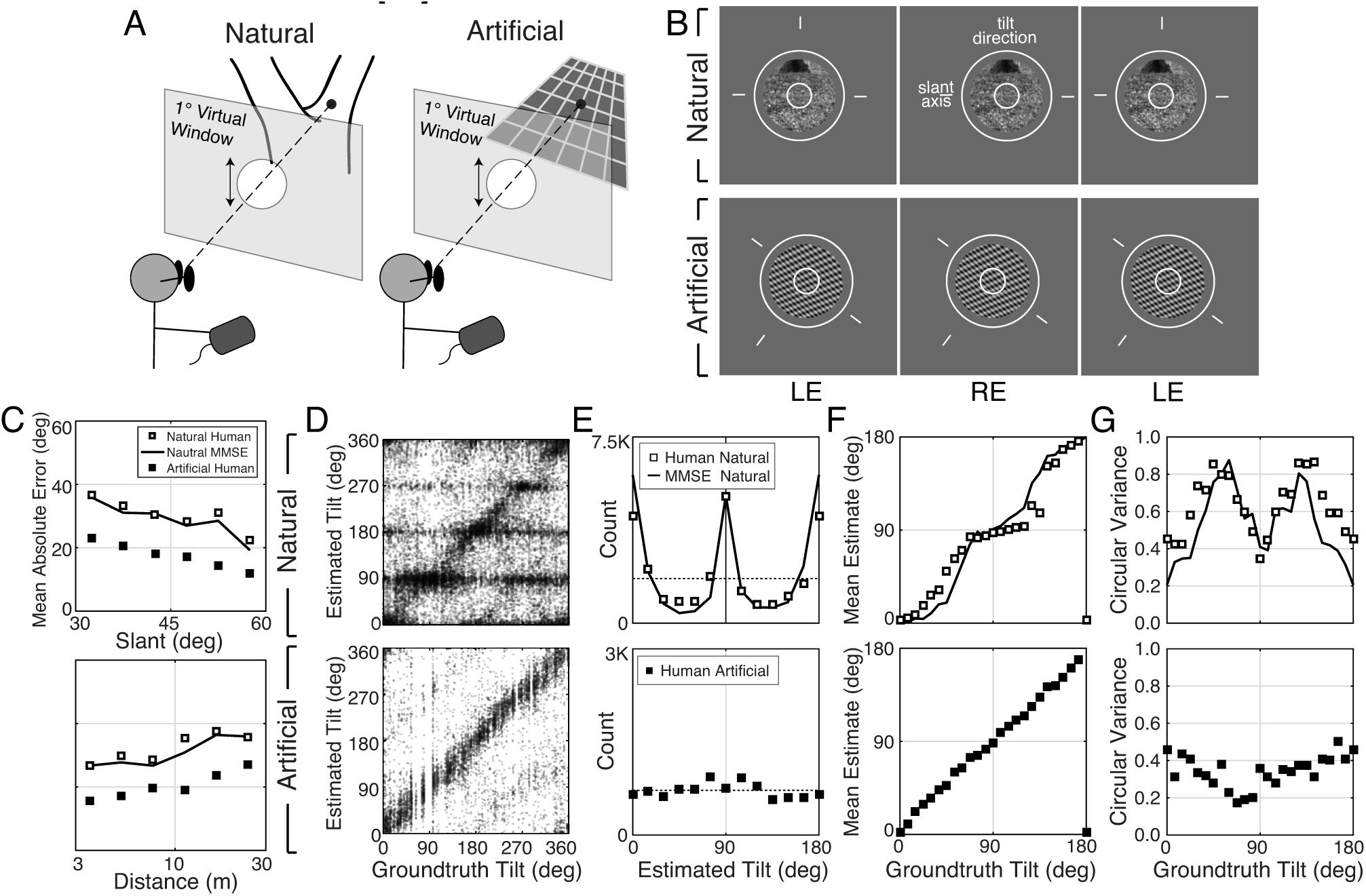
Experimental stimuli and human tilt responses. **A** The virtual viewing situation. **B** Natural and artificial stimuli (upper & lower rows). The task was to report the tilt at the center of the small (1° diameter) circle. Aperture sizes were either 3° (shown) or 1° (not shown). Observers set the orientation of the probe (circles and line segments) to indicate estimated tilt. Free-fuse to see in stereo 3D. **C** Impact of slant and distance on tilt estimation errors. **D** Raw responses for every trial in the experiment. **E** Histogram of raw responses (estimates) in the unsigned tilt domain (i.e. *τ* = [0°, 180°)). The dashed horizontal line shows the uniform distribution of groundtruth tilts presented in the experiment. (Histograms of signed tilt estimates are shown in Fig. S3.) **F** Mean estimates and **G** estimate variance as a function of groundtruth tilt. Human tilt estimates are more biased and variable with natural stimuli (top) than with artificial stimuli (bottom; 1/f noise, 3.5cpd plaid, and 5.25cpd plaid). With artificial stimuli, human estimates are unbiased and estimate variance is low. Model observer predictions (minimum mean squared error (MMSE) estimates; black curves) parallel human performance with natural stimuli.

Natural and artificial stimuli elicited strikingly different patterns of performance (Fig. 2D). Although many stimuli of both types elicit tilt estimates 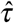 that approximately match the groundtruth tilt (i.e. data points on unity line), a substantial number of natural stimuli elicit estimates that cluster at the cardinal tilts (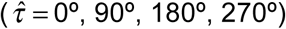). No such clustering occurs with artificial stimuli. A histogram of the human tilt estimates explicitly shows the clustering, or lack thereof (Fig. 2E). With natural stimuli, the distribution of estimates 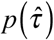 peaks at 0° and 90° and has a similar shape to the prior distribution of groundtruth tilts in the natural scene database (Fig. 1C; also see Fig. S3). If the database is representative of natural scenes, then one might expect the human visual system to use the natural statistics of tilt as a tilt prior in the perceptual processes that convert stimulus measurements into estimates. Standard Bayesian estimation theory predicts that the prior will influence estimates more when measurements are unreliable and influence estimates less when measurements are reliable [25].

We summarized 3D tilt estimation performance by computing the mean and variance of the tilt estimates 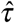as a function of groundtruth tilt (Fig. 2F,G). (The mean and variance were computed using circular statistics because tilt is an angular variable; see Methods.) These summary statistics change systematically with groundtruth tilt, exhibiting patterns reminiscent of the 2D oblique effect [26–28]. With natural stimuli, estimates are maximally biased at oblique tilts and unbiased at cardinal tilts; estimate variance is highest at oblique tilts (∼60° and ∼120°) and lowest at cardinal tilts. With artificial stimuli, estimates are essentially unbiased and are less variable across tilt. (The unbiased responses to artificial stimuli imply that the biased responses to natural stimuli accurately reflect biased perceptual estimates, under the assumption that the function mapping perceptual estimates to probe responses is stable across stimulus types; see Supplement.) The summary statistics reveal clear differences between the stimulus types. However, there is more to the data than the summary statistics can reveal. Thus, we analyzed the raw data more closely.

The probabilistic relationship between groundtruth tilt *τ* and human tilt estimates 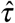 is shown in Figures 3 and 4. Each subplot in Fig. 3A shows the distribution of estimation errors 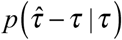 fora different groundtruth tilt. With artificial stimuli, estimation errors 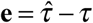 are unimodally distributed and peaked at zero (black symbols). With natural stimuli, estimation errors are more irregularly distributed, and the peak locations change systematically with groundtruth tilt (white points). Specifically, when the groundtruth tilt is a cardinal tilt (e.g. *τ* = 0° or *τ* = 90°), the error distributions peak at zero and large errors are rare. When the groundtruth tilt is an oblique tilt, the error distributions tend to be bi-modal with two prominent peaks at non-zero errors. Large errors are therefore more common with natural stimuli. For example, when groundtruth tilt *τ* = 60° the most common errors are −60° and 30°. These errors occur because observers incorrectly estimated the tilt to be 0° or 90°, respectively. Thus, with natural stimuli, human observers frequently (and incorrectly) estimate a cardinal tilt instead of the correct, oblique tilt.

**Figure 3.**
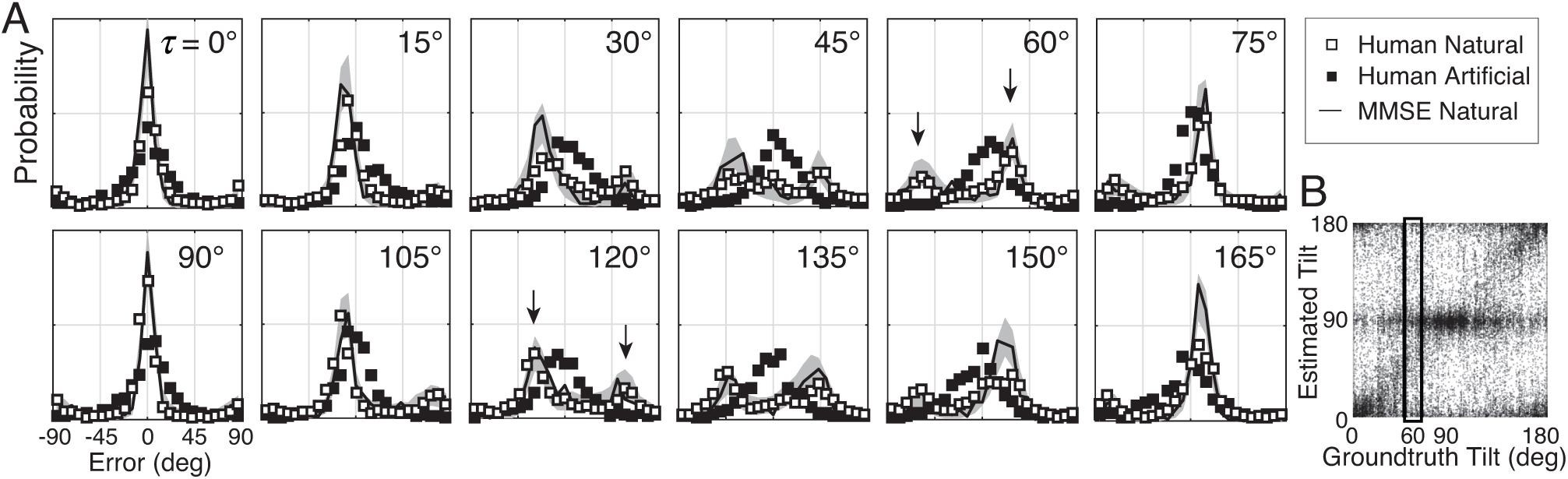
Distribution of tilt estimation errors for different groundtruth tilts. **A** Conditional error distributions 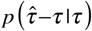 are obtained by binning estimates for each groundtruth tilt (vertical bins in B) and subtracting the groundtruth tilt. With artificial stimuli, the error distributions are centered on 0° (black symbols). With natural stimuli, the distributions of estimation error changes systematically with groundtruth tilt (white symbols). For cardinal groundtruth tilts (0° and 90°), the most common error is zero. For oblique tilts, the distribution of tilt errors with natural stimuli peaks at values other than zero (e.g. arrows in *τ* = 60° and *τ* = 120° subplots). The irregular error distributions are nicely predicted by the MMSE estimator (black curve); shaded regions show 95% confidence intervals on the MMSE estimates from 1000 Monte Carlo simulations of the experiment. The MMSE estimator had zero free parameters to fit to the human responses. **B** Raw unsigned tilt estimates with natural stimuli. The box shows estimates in the *τ* = 60° tilt bin. This data is the same as in Fig. 2D, but is shown in the unsigned tilt domain.

**Figure 4.**
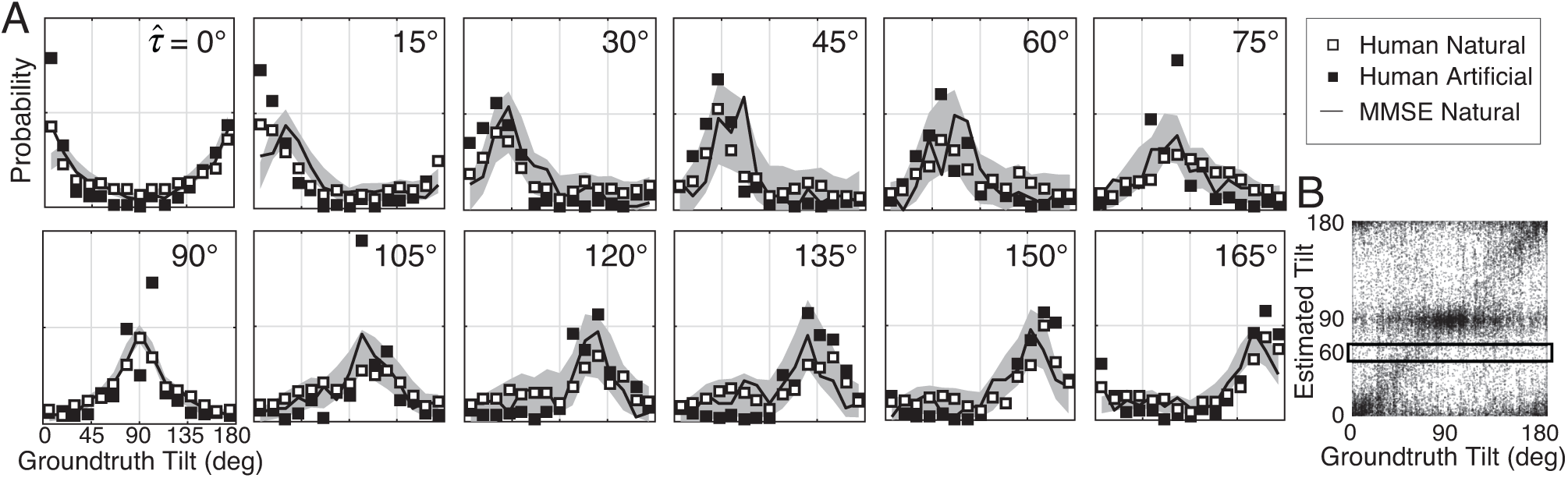
Distribution of groundtruth tilts for different tilt estimates. **A** Conditional distributions of groundtruth tilt 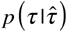) are obtained by binning groundtruth tilts for each estimated tilt (horizontal bins in B). Unlike the conditional error distributions, these distributions with natural and artificial stimuli are similar. The most probable groundtruth tilt, conditional on the estimate, peaks at the estimated tilt for both stimulus types. Thus, any given estimate is a good indicator of the groundtruth tilt despite the overall poor performance with natural stimuli. Also, these conditional distributions are well accounted for by the MMSE estimates; shaded regions show 95% confidence intervals on the MMSE estimates from 1000 Monte Carlo simulations of the experiment. The MMSE had zero free parameters to fit to human performance. **B** Raw unsigned tilt estimates. The box shows groundtruth tilts in the 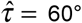 estimated tilt bin.

Tilt estimates from natural stimuli are less accurate at oblique than at cardinal groundtruth tilts. Does this fact imply that oblique tilt estimates (e.g. 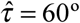) provide less accurate information about groundtruth tilt than cardinal tilt estimates (e.g. 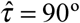)? No. Each panel in Fig. 4A shows the distribution of groundtruth tilts 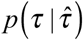 for each estimated tilt. The most probable groundtruth tilt equals the estimated tilt, and the variance of each distribution is approximately constant, regardless of whether the estimated tilt is cardinal or oblique. Furthermore, the estimates from natural and artificial stimuli provide nearly equivalent information about groundtruth (see also Fig. S4). Thus, even though tilt estimation performance is far poorer at oblique than at cardinal tilts, and is far poorer in natural than in artificial stimuli, all tilt estimates regardless of value are similarly good predictors of groundtruth tilt.

How can it be that low-accuracy estimates from natural stimuli predict groundtruth nearly as well as high-accuracy estimates from artificial stimuli? Some regions of natural scenes yield high-reliability measurements that make tilt estimation easy; other regions of natural scenes yield low-reliability measurements that make tilt estimation hard. When measurements are reliable, the prior influences estimates less; when measurements are unreliable, the prior influences estimates more. Thus, cardinal tilt estimates can result either from reliable measurements of cardinal tilts or from unreliable measurements of oblique tilts. On the other hand, oblique tilt estimates can only result from reliable measurements of oblique tilts, because the measurements must be reliable enough to overcome the influence of the prior. All these factors combine to make each tilt estimate, regardless of its value, an equally reliable indicator of groundtruth tilt. The uniformly reliable information provided by the estimates about groundtruth (see Fig. 4A) may simplify the computational processes that optimally pool local into global estimates (see Discussion). The generality of this phenomenon across natural tasks remains to be determined. However, we speculate that it may have widespread importance for understanding perception in natural scenes, and in other circumstances where measurement reliability varies drastically across spatial location.

### Normative model

We asked whether the complicated pattern of human performance with natural stimuli is consistent with optimal information processing. To answer this question, we compared human performance to the performance of a normative model, a Bayes optimal observer that optimizes 3D tilt estimation in natural scenes given a squared error cost function [21]. The model takes three local image cues **C** as input—luminance, texture, and disparity gradients—and returns the minimum mean squared error (MMSE) tilt estimate 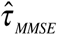 as output. (The MMSE estimate is the mean of the posterior probability distribution over groundtruth tilt given the measured image cues.)

To determine the optimal estimate for each possible triplet of cue values, we use the natural scene database. At each pixel in the database, the image cues are computed directly from the photographic images within a local area, and the groundtruth tilt is computed directly from the distance data (see Methods; [21]). In other words, the model is ‘image-computable’: the model computes the image cues from image pixels and produces tilt estimates as outputs.

We approximate the posterior mean *E*[*τ* | **C]** = ∑*_τ_ τ p*(*τ* | **C**) by computing the sample mean of the groundtruth tilt conditional on each unique image cue triplet (Fig. 5A). The result is a table, or ‘estimate cube’, where each cell stores the optimal estimate 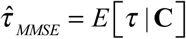 for a particular combination of image cues (Fig. 5B).

**Figure 5.**
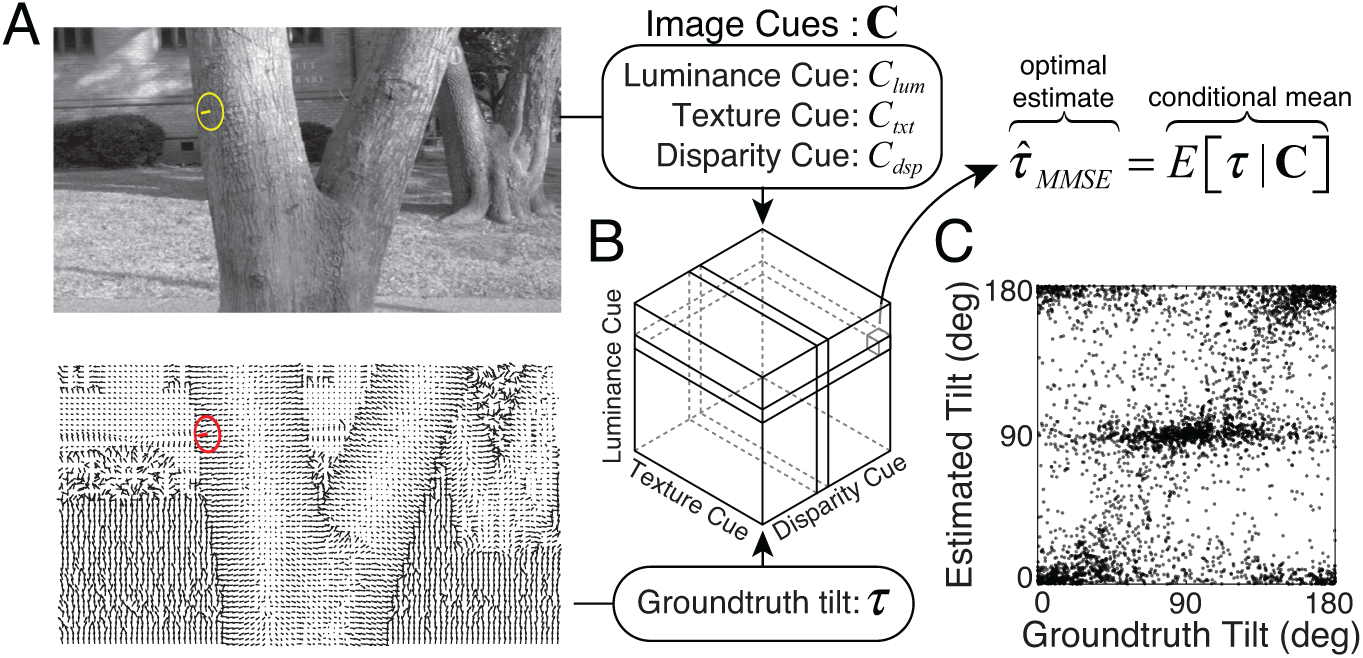
Normative model for tilt estimation in natural scenes. **A** The model observer estimate is the minimum mean squared error (MMSE) tilt estimate 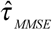 given three image cue measurements. Optimal estimates are approximated from 600 million data points (90 stereo-images) in the natural scene database: image cue values are computed directly from the photographic images and groundtruth tilts are computed directly from the distance data. **B** MMSE estimates for ∼260,000 (64^3^) unique image cue triplets are stored in an ‘estimate cube’. **C** Model observer estimates for the 3600 unique natural stimuli used in the experiment. For each stimulus used in the experiment, the image cues are computed, and the MMSE estimate is looked up in the ‘estimate cube’. The optimal estimates within the estimate cube change smoothly with the image cue values; hence, a relatively small number of samples can explore the structure of the full 3D space and provide representative performance measures (see Discussion).

In the cue-combination literature, cues are commonly assumed to be statistically independent [29]. In natural scenes, it is not clear whether this assumption holds. Fortunately, the current model is free of assumptions about statistical independence and the form of the joint probability distribution (see Discussion). Thus, our normative model provides a principled benchmark, grounded in natural scene statistics, against which to compare human performance.

We tested the model observer on the exact same set of natural stimuli used to test human observers (Fig. 5C). The model observer predicts the overall pattern of raw human responses (Fig. S5). More impressively, the model observer also predicts the mean and variance of the human tilt estimates (Fig. 2D-G), the conditional error distributions (Fig. 3), and the conditional groundtruth tilt distributions (Fig. 4). We conclude that the biased and imprecise human tilt estimates with natural stimuli are nevertheless lawful.

To ensure that the predictive power of the MMSE estimator is not trivial, we developed multiple alternative model observers. None of the other model observers predict human performance as well as the MMSE estimator (Fig. S6). Our results do not rule out the possibility that another model could predict human performance better, but the current MMSE estimator establishes a strong benchmark against which other models must be compared.

Two points are worth emphasizing. First, this model observer had no free parameters that were fit to the human data [21]; instead, the model observer was designed to perform the task optimally given the three image cues. Second, the close agreement between human and model performance suggests that humans use the same cues (or cues that strongly correlate with those) used by the normative model (see Discussion).

### Trial-by-trial Error

If human and model observers use the same cues in natural stimuli to estimate tilt, variation in the stimuli should cause similar variation in performance. Is human and model observer performance similar on individual trials? The same set of natural stimuli was presented to all observers. Thus, it is possible to make direct, trial-by-trial comparisons of the estimation errors that each observer made. If the properties of individual natural stimuli influence estimates similarly across observers, then observer errors across trials should be correlated. Accounting for trial-by-trial errors is one of the most stringent comparisons that can be made between a model and human performance.

Natural stimuli indeed elicit similar trial-by-trial errors from human and model observers (Fig. 6A). The model predicts trial-by-trial human errors far better than chance. We quantify the model-human similarity with the circular correlation coefficients of the trial-by-trial model and human estimates (Fig. 6B). The correlation coefficients are significant. This result implies that the errors are systematically and reliably dependent on the properties of natural stimuli, and that these properties affect human and model observers similarly.

**Figure 6.**
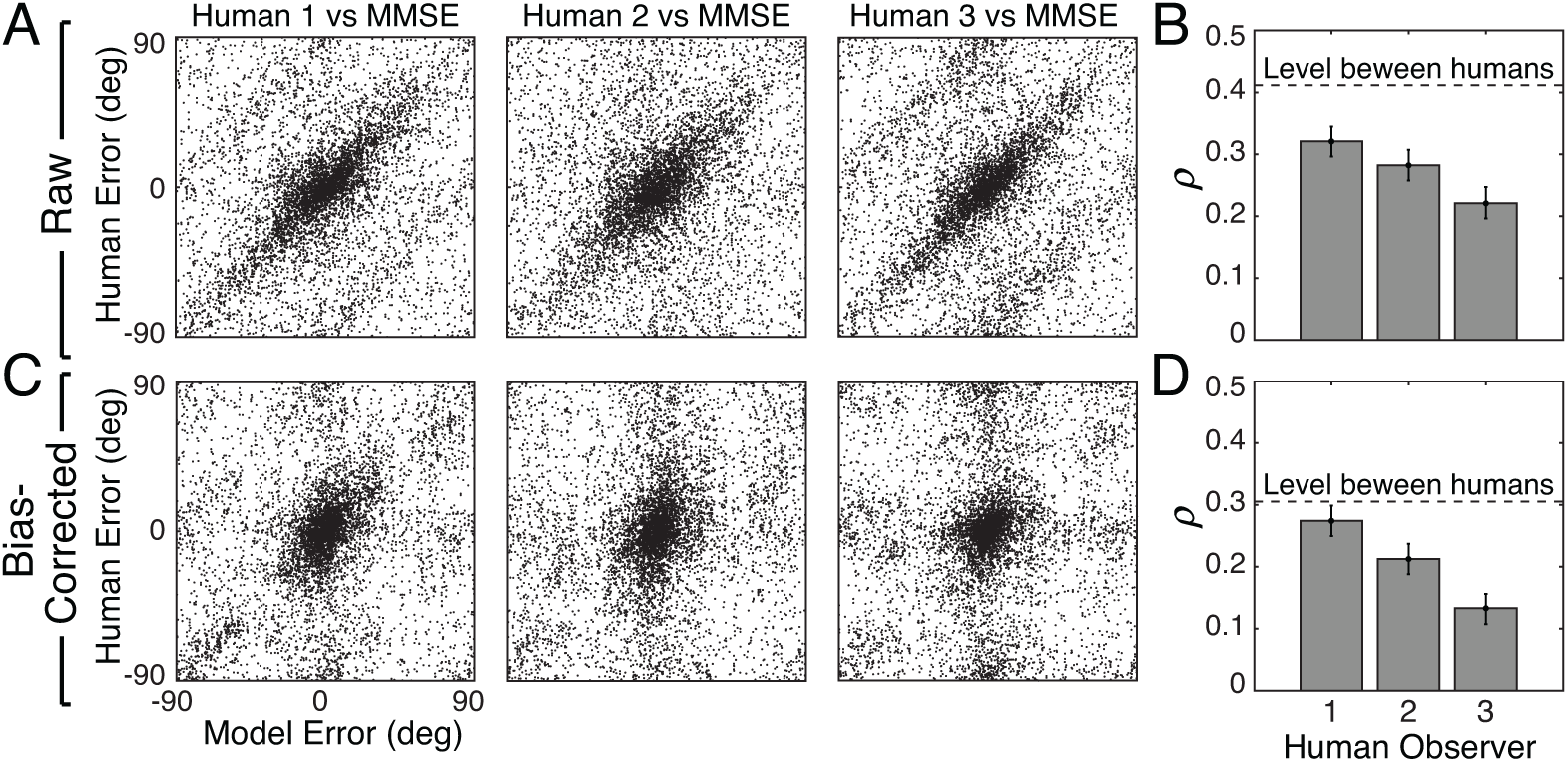
Trial-by-trial estimation errors: model vs. human observers. The diagonal structure in the plots indicates that trial-by-trial errors are correlated. **A** Raw trial-by-trial errors with natural stimuli between model and human observers. **B** Correlation coefficients (circular) for trial-by-trial errors between model and each human observer. The error bars represent 95% confidence intervals from 1000 bootstrapped samples of the correlation coefficient. The dashed line shows the mean of correlation coefficients of errors between human observers in natural stimuli (Fig. S7). **C** Bias-corrected errors in natural stimuli. **D** Correlation coefficient for bias-corrected errors.

However, because both human and model observers produced biased estimates with natural stimuli (Fig. 2F, Fig. S2), it is possible that the biases are responsible for the error correlations. Thus, we must remove the influence of bias. To do so, we compute the bias-corrected error. On each trial, the observer bias at each groundtruth tilt was subtracted 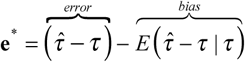 rom the raw error. The bias-corrected errors of the human and model observers also significantly correlated (Fig. 6C,D). The human-human correlation (dashed line, Fig. 6B,D; Fig. S7) sets an upper bound for the model-human correlation. The model-human correlation approaches this bound for some humans. Other measures of trial-by-trial similarity (e.g., choice probability; Fig. S6) yield similar conclusions. These results show that natural stimulus variation at a given groundtruth tilt causes similar response variation in human and model observers.

Thus, the normative model, which was not fit to the human data, accounts for human tilt estimates at the level of the summary statistics (Fig. 2E-G), the conditional distributions (Figs. 3,4), and the trial-by-trial errors (Fig. 6). Together, this evidence suggests that the human visual system’s perceptual processes and the normative model’s computations are making similar use of similar information. We conclude that the human visual system makes near-optimal use of the available information in natural stimuli for estimating 3D surface tilt.

### Effect of Natural Depth Variation

Natural and artificial stimuli were matched on many dimensions: tilt, slant, distance, and luminance contrast. But tilt estimation with natural stimuli was considerably poorer than tilt estimation with artificial stimuli (Fig. 2,3). Why? Other factors must account for the performance differences. Natural scenes contain natural depth variation (i.e. complex surface structure); some surfaces are approximately planar, some are curved or bumpy. In our experiment, each artificial scene consisted of one planar surface. We checked whether performance differences between natural and artificial scenes can be attributed to differences in surface planarity. We quantified the departure of surface structure from planarity by computing local *tilt contrast power* (i.e. tilt variance; see Methods) in the central 1° area of each natural stimulus. Then, we examined how estimation error changes with tilt contrast power.

Human and model observer errors increase linearly with tilt contrast power (Fig. 7A); at the smallest tilt contrasts (near-planar natural stimuli), overall human errors with natural and artificial stimuli are nearly identical. However, tilt contrast co-varies with groundtruth tilt: cardinal tilts tend to have lower tilt contrast than oblique tilts (Fig. S8) presumably because of the ground plane. This means that we could be misattributing the effect of groundtruth tilt to tilt contrast. Therefore, we repeated the analysis for cardinal tilts and oblique tilts separately. We found that tilt contrast continues to have a large effect on performance (Fig. 7B). Thus, like slant and distance (Fig. 2C), we have shown that tilt contrast (surface planarity) is a key stimulus dimension that strongly influences tilt estimation performance.

**Figure 7.**
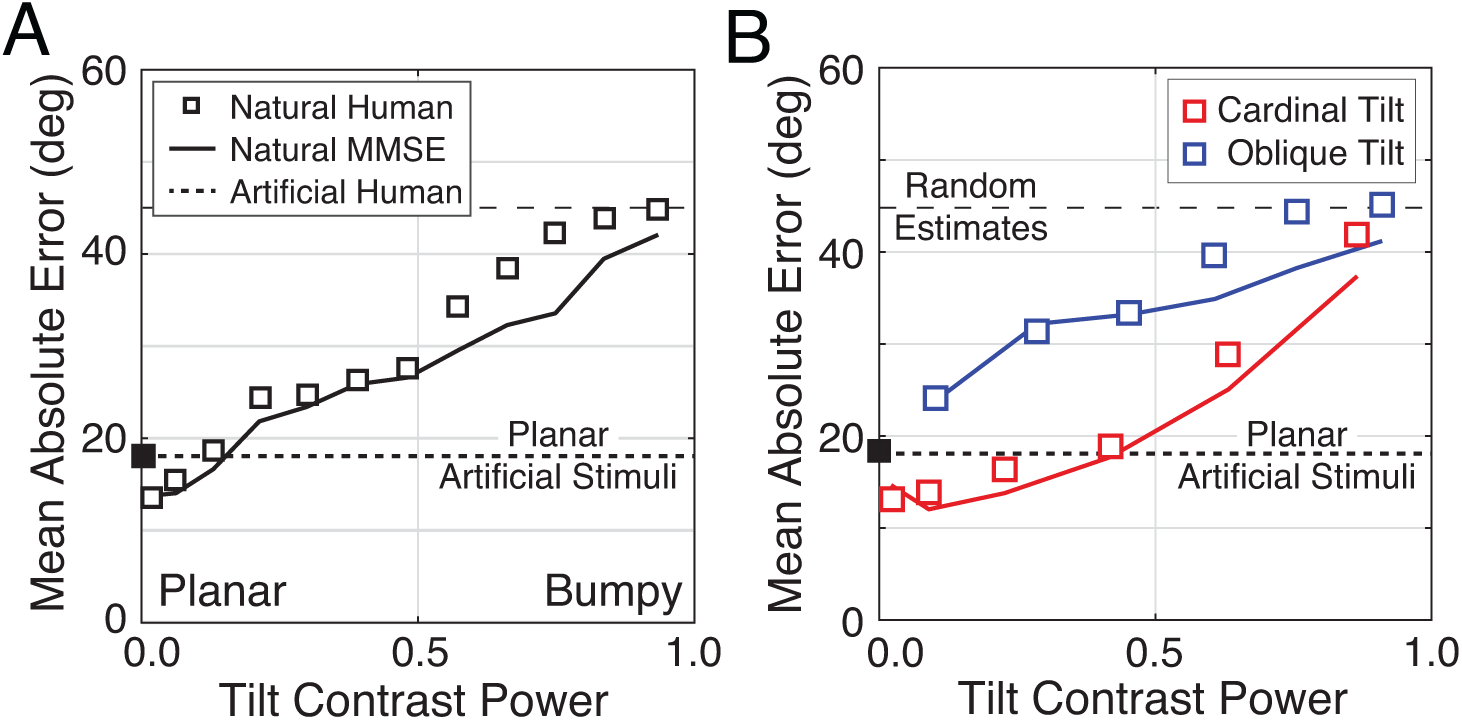
The effect of local depth variation on tilt estimation error. **A** Absolute error increases linearly with tilt contrast power (i.e. tilt variance) in natural stimuli for humans and model observers. Artificial stimuli were perfectly planar and had zero local depth variation; hence the individual data point at zero tilt contrast power. **B** Same as A, but conditional on whether the groundtruth tilts are cardinal (red) or oblique (blue). Data points are spaced unevenly because they are grouped in quantile bins, such that each data point represents an equal number of stimuli. The solid curves represent the errors of the MMSE estimator for cardinal (red) and oblique (blue) groundtruth tilts. The normative model predicts performance in all cases.

The effect of tilt contrast on tilt estimation is similar to the effect of luminance contrast on target detection, a classic task in the spatial vision literature. In target detection tasks, for example, threshold contrast power increases linearly with noise contrast power [30], just as tilt estimation error increases linearly with tilt contrast power. We speculate that characterizing the impact of tilt contrast will be as important to understanding human 3D surface orientation estimation as modeling local luminance contrast has been to understanding how humans detect spatial patterns in images.

Do differences in tilt contrast account for *all* performance differences between natural and commonly used artificial stimuli? No. Human tilt estimates of near-planar surfaces in natural scenes still tend to be more biased and exhibit a different pattern of precision than tilt estimates from the artificially stimuli used in our experiments (Fig. S9). Understanding the factors that contribute to these remaining differences is an important direction for future work.

## Discussion

3D surface orientation estimation is a classic task in vision science, requiring the estimation of slant and tilt. The current study focuses on tilt estimation. We quantify performance in natural scenes and report that human tilt percepts are often neither accurate nor precise. To connect our work to the classic literature, we matched artificial stimuli to the natural stimuli on the stimulus dimensions that are controlled in typical experiments. The comparison revealed dramatic performance differences. Furthermore, the detailed patterns of human performance are predicted without free parameters by a normative model, grounded in natural scene statistics, that makes the best possible use of available image information. Importantly, the normative model is distinguished from many models of mid-level visual tasks because it is ‘image computable’; that is, it takes image pixels as inputs and produces tilt estimates as outputs. Together, the current experiment and modeling effort contributes to a broad goal in vision and visual neuroscience research: to generalize our understanding of human vision from the lab to the real world.

### Generality of Conclusions and Future Directions

#### Influence of scale

Groundtruth surface orientation is computed from a locally planar approximation to the surface structure, but surfaces in natural scenes are generally non-planar. Hence, the area over which groundtruth tilt is computed can affect the values assigned to each surface location. The same is true of the local image cue values. We checked how sensitive our results our to the scale of the local analysis area. We recomputed groundtruth tilt for two scales and recomputed image cue values for four scales (see Methods). All eight combinations of scales yield the same qualitative pattern of results.

#### Influence of gaze angle

The statistics of local surface orientation change with elevation in natural scenes [23,31]. In our study, scene statistics were computed from range scans and stereo-images (36°x21° field-of-view) that were captured from human eye height with earth parallel gaze [21]. Different results may characterize other viewing situations, a possibility that could be evaluated in future work. However, the vast majority of eye movements in natural scenes are smaller than 10° [32–34]. Hence, the results presented here are likely to be representative of an important subset of conditions that occur in natural viewing.

#### Influence of internal noise

The normative model (i.e. MMSE estimator) used in this paper does not explicitly model the effect of internal noise on task performance. However, it may be that in natural scenes, natural stimulus variation is the controlling source of noise. If so, the explicit modeling of noise may be unnecessary. Determining the relative importance of natural stimulus variability and internal noise is an important topic for future work.

#### Influence of sampling error

The natural stimuli presented in the experiment were chosen via constrained random sampling (see Methods). Random stimulus sampling increases the likelihood that the reported performance levels are representative of generic natural scenes. One potential concern is that the relatively small number of unique stimuli that can be practically used in an experiment (e.g. n=3600 in this experiment) precludes a full exploration of the space of optimal estimates (see Fig. 5B). Fortunately, the tilt estimates from the normative model change smoothly with the image cue values (14). Systematic sparse sampling should thus be sufficient to explore the space. To rigorously determine the influence of each cue on performance, future parametric studies should focus on the role of particular image cue combinations and other important stimulus dimensions like tilt contrast.

#### Influence of non-optimal cues

Although the cues used by the normative model are widely studied and commonly manipulated, there is no guarantee that they are the most informative cues in natural scenes. Automatic techniques could be used to find the most informative cues for the task [35,36]. These techniques have proven useful for other visual estimation tasks with natural stimuli [37–40]. However, in the current task, we speculate that different local cues are unlikely to yield substantially better performance [21]. Also, given the similarities between human and model observer performance, any improved ability to predict human performance is likely to be modest at best. Nevertheless, the only way to be certain is to check, which we plan to do in the future.

#### 3D surface orientation estimation

The estimation of the three-dimensional structure of the environment is aided by the joint estimation of tilt and slant (Marr’s “2.5D sketch”) [41]. Although we have shown that human and model tilt estimation performance is systematically affected by surface slant (Fig. 2C), the current experiment and modeling work only address the human ability to estimate tilt. We have not yet explicitly tested or modeled how humans estimate slant or jointly estimate slant and tilt. In the future, we will extend the modeling framework to predict the human ability to jointly estimate tilt and slant (i.e. 3D surface orientation) in natural scenes.

### Cue-combination with and without independence assumptions

The standard approach to modeling cue-combination, sometimes known as maximum likelihood estimation, includes a number of assumptions: a squared error cost function, cue independence, unbiased Gaussian-distributed single cue estimates, and a flat or uninformative prior [29] (but see [42]). The approach used here (normative model; see Fig. 5) assumes only a squared error cost function, and is guaranteed to produce the Bayes optimal estimate given the image cues, regardless of whether common assumptions hold. In natural scenes, it is often unclear whether common assumptions hold. Methods with relatively few assumptions can therefore be powerful tools for establishing principled predictions. We have not yet fully investigated how image cues are combined in tilt estimation, but we have conducted some preliminarily analyses. For example, a simple average of the single-cue estimates (each based on luminance, texture, or disparity alone) underperforms the three-cue normative model used throughout the paper (see Fig. S6). This result is not surprising given that the individual cues are not independent, that the single cue estimates are not Gaussian distributed, and that the tilt prior is not flat. However, the current study is not specifically designed to examine the details of cue combination in tilt estimation. To rigorously examine cue-combination in this task, a parametric stimulus-sampling paradigm should be employed, a topic that will be explored in future work.

### Local and global estimation of tilt

A grand problem in perception and neuroscience research is to understand how local estimates are grouped into more accurate global estimates. We showed that that local tilt estimates are unbiased predictors of groundtruth tilt and have nearly equal reliability (Fig. 4). This result implies that optimal spatial pooling of the local estimates may be relatively simple; assuming statistical independence (i.e. naïve Bayes), optimal spatial pooling is identical to a simple linear combination of the local estimates: the straight average of *N* local estimates 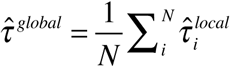. Of course, local groundtruth tilts and estimates are spatially correlated, so the independence assumption will not be strictly correct. However, the spatial correlations could be estimated from the database and incorporated into the computations [42]. Our work thus lays a strong empirically grounded foundation for the investigation of local-global processing in surface orientation estimation.

### Behavioral experiments with natural images

In classic studies of surface orientation perception, stimuli are usually limited in at least one of two important respects. If the stimuli are artificial (e.g. computer-graphics generated), groundtruth surface orientation is known but lighting conditions and textures are artificial, and it is uncertain whether results obtained with artificial stimuli will generalize to natural stimuli. If the stimuli are natural (e.g. photographs of real scenes), groundtruth surface orientation is typically unknown which complicates the evaluation of the results. The experiments reported here use natural stereo-images with laser-based measurements of groundtruth surface orientation, and artificial stimuli matched to the tilt, slant, distance, and contrast of the natural stimuli. This novel design allows us to relate our results to the classic literature, determine the generality of results with both natural and artificial stimuli, and isolate performance-controlling differences between the stimuli.

### Perception and the internalization of natural scene statistics

The current study is the latest in a series of reports that have attempted, with ever increasing rigor, to link properties of perception to the statistics of natural images and scenes. Our contribution extends previous work in several respects. First, previous work demonstrated similarity between human and model performance only at the level of summary statistics [28,43-45]. We demonstrate that a principled model, operating directly on image data, predicts the summary statistics, the distribution of estimates, and the trial-by-trial errors. Second, previous work showed that human observers behave as if their visual systems have encoded the task-relevant statistics of 2D natural images [28]. We show that human observers behave as if they have properly encoded the task-relevant joint statistics of 2D natural images and 3D properties of natural scenes (also see [43]. Third, previous work tested and modeled human performance with artificial stimuli only [28,43–45]. We test human performance with both natural and artificial stimuli. The dramatic, but lawful, fall-off in performance with natural stimuli highlights the importance of performing studies with the stimuli visual systems evolved to process.

## Author Contributions

SK and JB conceived the project, designed the experiments, carried out the data analysis and computational modeling, and wrote the manuscript.

## Methods

### Apparatus

The stereo images were presented with a ViewPixx Technologies ProPixx projector fitted with a 3D polarization filter. Left and right images were presented sequentially at a refresh rate of 120 Hz (60 Hz per eye) with the same resolution of the images (1920x1080 pixel). The observer was positioned 3.0m from a 2.0x1.2m Harkness Clarus 140 XC polarization maintaining projection screen. This viewing distance minimizes the potential influence of screen cues to flatness (e.g. blur). Human observers wore glasses with passive (linear) polarized filters to isolate the image for the left and right eyes. The observer’s head was stabilized with a chin- and forehead-rest. From this viewing position, the projection screen subtended 36°x21° of visual angle. The disparity-specified distance created by this projection system matched to the distances measured in the original natural scenes. The projection display was linearized over 10bits of gray level. The maximum luminance was 84 cd/m^2^. The mean luminance was set to 40% of the projection system’s maximum luminance.

### Experiment

Three human observers, the two authors and one naïve subject, binocularly viewed a small region of a natural scene through a circular aperture (1° or 3° diameter) positioned 5 arcmin of disparity in front of the scene point. Observers communicated their tilt estimate with a mouse-controlled probe. Each observer viewed 3600 unique natural stimuli (150 stimuli per tilt bin x 24 tilt bins) presented with each of two apertures in the experiment (7200 total). Natural stimuli were constrained to be binocularly visible (no half-occlusions), have slants larger than 30°, have distances between 5m and 50m, and have contrasts between 5% and 40%. Each observer also viewed 1440 unique artificial stimuli (60 stimuli per tilt bin x 24 tilt bins) with two apertures (2880 total). Artificial stimuli (1/f noise and phase- and orientation-randomized plaids) were matched to the natural stimuli on multiple additional dimensions (tilt, slant, distance, and contrast). Natural stimuli were presented in 48 blocks of 150 trials each, and artificial stimuli data were presented in 12 blocks of 240 trials each, with interleaved blocks using small and large apertures.

### Data analysis

Tilt is a circular (angular) variable. We computed the mean, variance, and error using standard circular statistics. The circular mean is defined as 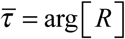 where 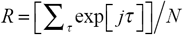 is the complex mean resultant vector. The variance is defined as var (*τ*) = 1 − |*R*|. Estimation error 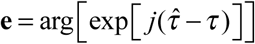 is the circular distance between the tilt estimate and groundtruth.

### Groundtruth tilt

Groundtruth tilt *τ* is computed from the distance data (range map **r**) co-registered to each natural image in the database. We defined groundtruth tilt *τ* = atan2(∇ _*y*_**r**,∇_*x*_**r**) as the orientation of the normalized range gradient [41]. The range gradient was computed by convolving the groundtruth distance data with a 2D Gaussian having space constant *σ* and then taking the partial derivatives in *x* and *y* directions on the image [21]. For the results presented in this manuscript, groundtruth tilt was computed using a space constant of *σ* = 3 arcmin (patch size at half height ≅ 0.1°); doubling this space constant (patch size ≅ 0.25°) does not change the qualitative results.

### Image cues to tilt

Image cues to tilt (disparity, luminance, and texture) were computed directly from the images. Like groundtruth tilt, disparity and luminance cues to tilt are defined as the orientation *C* = atan2(∇ _*y*_*cue*,∇_*x*_ *cue*) of the local disparity and luminance gradients. The local disparity gradient is computed on the disparity image, which is obtained from the left and right eye luminance images via standard local windowed cross-correlation [21,46,47]. The texture cue to tilt is defined as the orientation of the major axis of the local amplitude spectrum. This texture cue is non-standard (but see [8]). However, we have previously verified that this texture cue is more accurate in natural scenes than more traditional texture cues [21,48–51]. For the main results presented in this manuscript, image cues were computed using a space constant of *σ* = 6 arcmin (patch size at half-height ≅ 0.25°); changing the space constants to *σ* = 3, 6, 9, or 12 arcmin (patch size at half-height ≅ 0.1-0.5°) does not change the qualitative results.

### Local tilt contrast power

Each pixel under an aperture has a corresponding groundtruth tilt. The groundtruth tilt variance, which is referred to as the tilt contrast power in the main text, is the circular variance of the tilt values under a 1° aperture.

## Supplement

### The output-mapping problem

On each trial, human observers communicated their perceptual estimate 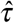 by making a response 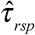 with a mouse-controlled probe. Unfortunately, the responses are not guaranteed to equal the perceptual estimates. An output-mapping function 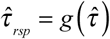 relates the response to the perceptual estimate, and an estimation function 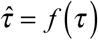 relates the estimate to the groundtruth tilt of each stimulus. When responses are biased, it is hard to conclude whether the biases are due to the output-mapping function or to the estimation function. When responses are unbiased, a stronger case can be made that the human responses equal the perceptual estimates. To obtain unbiased responses 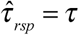 from biased estimates 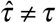, the output mapping function would have to exactly equal the inverse of a biased estimation function: *g* (.) = *f* ^−1^(.); this possibility seems unlikely and has no explanatory power. Thus, by Occam’s razor, unbiased responses imply unbiased output-mapping and estimation functions: 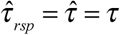. Human responses to artificial stimuli were unbiased (Fig. 2F), implying an unbiased output-mapping function. Assuming that the output-mapping function is stable across stimulus types, we conclude that the biased-observer responses to natural stimuli accurately reflect biased perceptual estimates.

**Figure S1.**
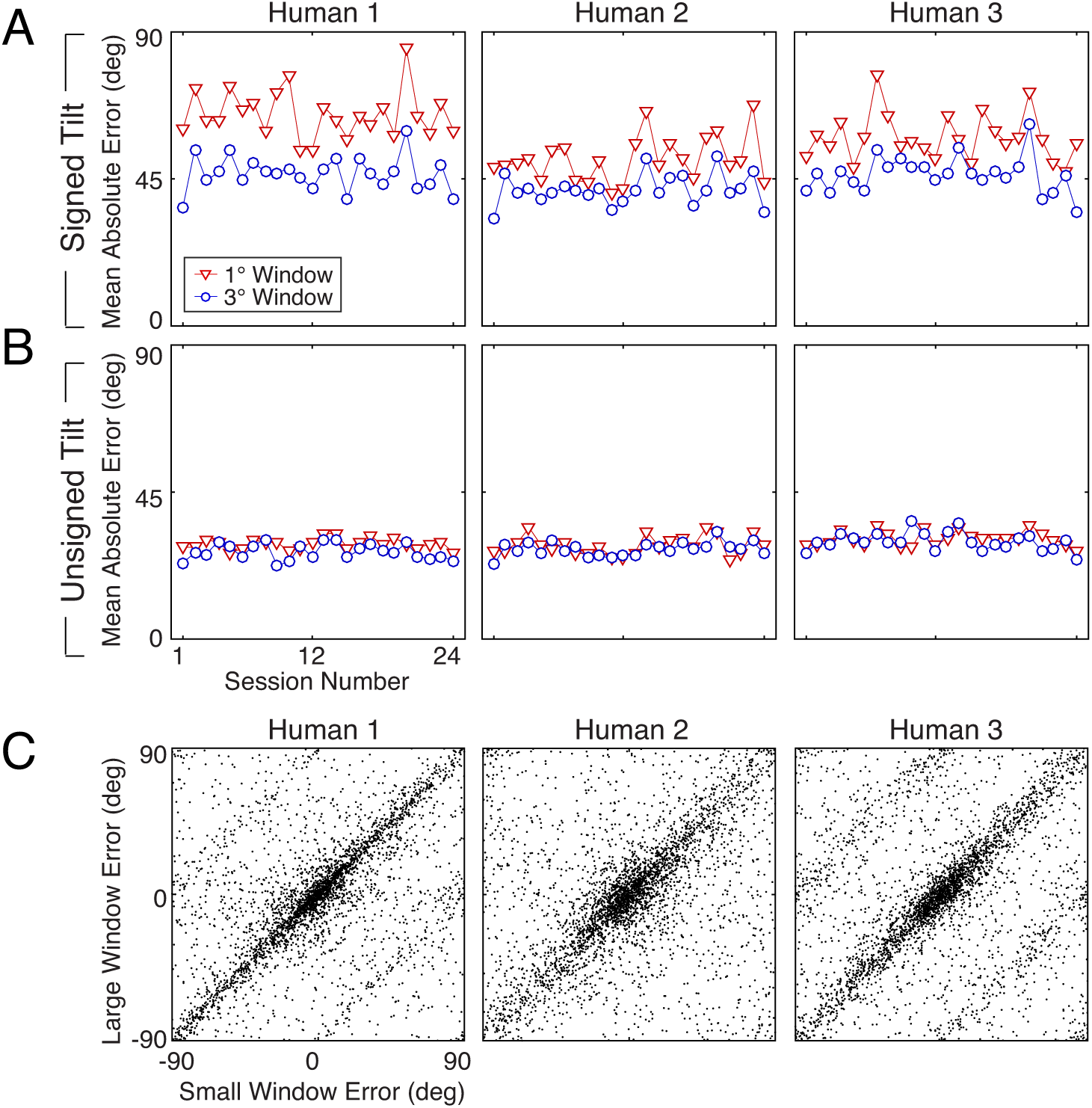
Tilt estimation errors with small (1°, red) vs. large (3°, blue) apertures for natural stimuli. Signed tilt is defined in *τ* = [0°,360°) with slant in [0°,90°); unsigned tilt is defined in *τ* = [0°,180°) and slant in [-90°,90°). The errors are analyzed with signed or unsigned domain. To reduce the possibility that individual stimuli were memorized, sessions with small apertures always preceded sessions with large apertures for a given set of natural stimuli. **A** Mean absolute error in the signed tilt domain across trials in each of 24 experimental sessions. The 3° aperture benefits signed tilt estimation performance. With a 1° aperture, humans make significantly more sign mistakes (e.g. estimating a tilt of 45° when the groundtruth tilt is 225°). **B** Mean absolute error in the unsigned tilt domain across trials in each of 24 experimental blocks. Errors with large and small apertures in the unsigned domain are nearly indistinguishable. **C** Unsigned trial-by-trial errors with small vs. large windows for each human observer. The errors are strongly correlated, although there are several examples of 90° shifted estimates in the data of Humans 1 and 3. Overall, with unsigned tilt, performance with small and large windows was nearly equivalent.

**Figure S2.**
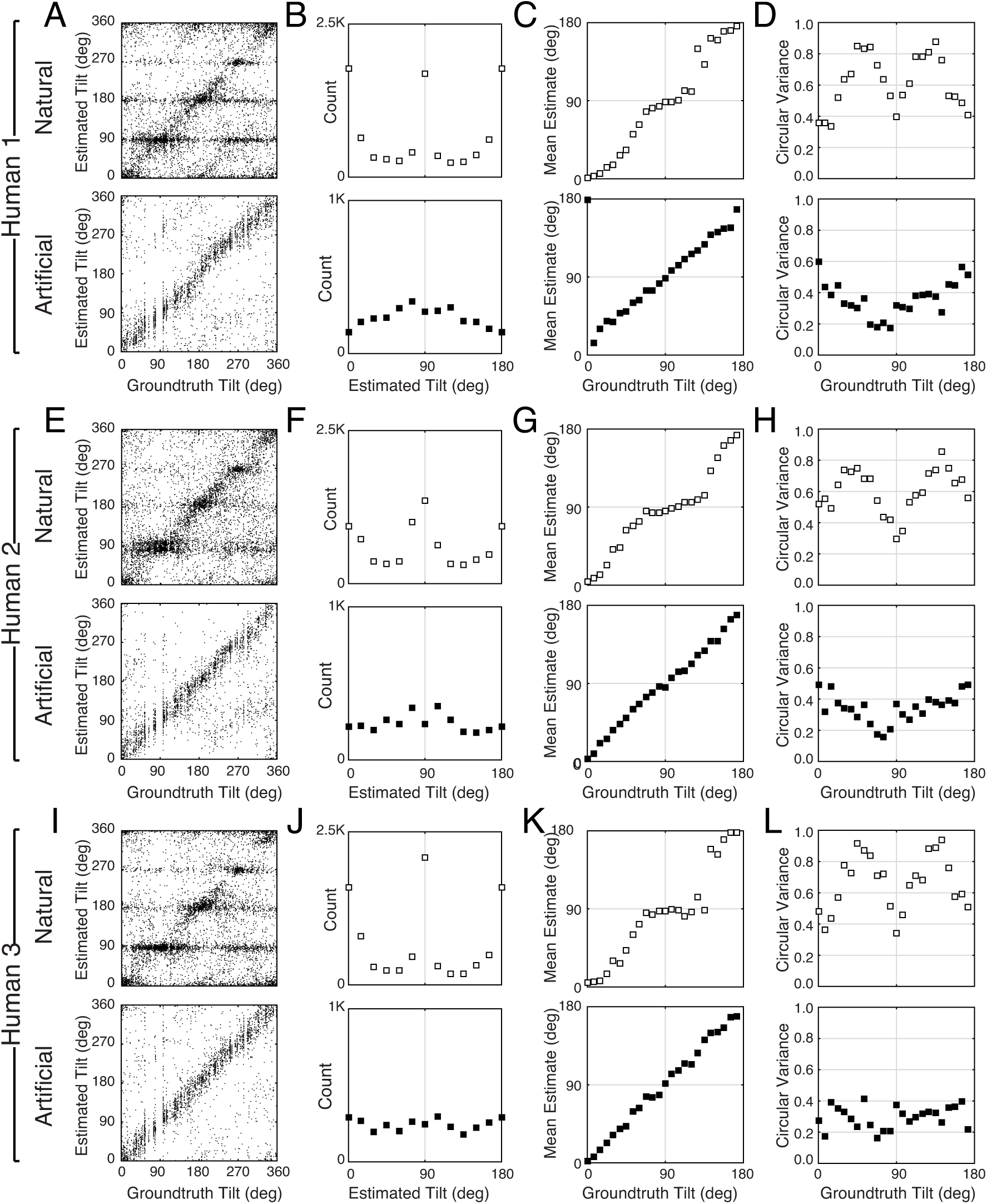
Tilt estimation performance for individual human observers. **A** Human 1 raw tilt responses, represented in the signed tilt domain *τ* = [0° 360°), for natural and artificial stimuli. **B** Histogram of raw responses (estimates) in the unsigned domain *τ* = [0° 180°). **C** Mean estimates and **D** estimate variance as a function of groundtruth tilt. **E-H** Same as A-D but for Human 2. **I-L** Same as A-D but for Human 3.

**Figure S3.**
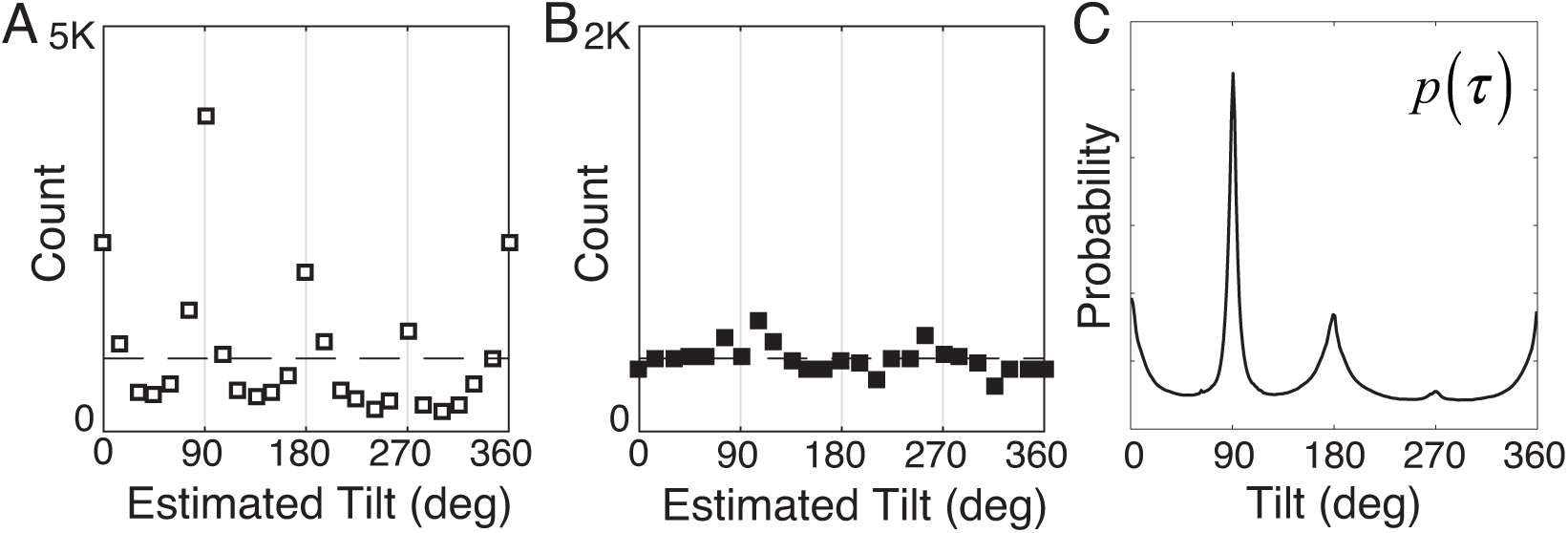
Histogram of raw responses (estimates) from human observers in signed tilt domain *τ* = [0° 360°). The dashed line represents the uniform distribution of tested stimuli. **A** Human responses to natural stimuli. The distribution 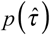 peaks at cardinal angles, but the peak at 270° is significantly lower than the peak at 90°, similar to the distribution of tilts in natural scenes *p*(*τ*) computed directly from the database. **B** Human responses to artificial stimuli are more similar to the distribution of tested tilts. **C** Prior probability distribution of groundtruth tilts, computed directly from the natural scene database, in the signed tilt domain.

**Figure S4.**
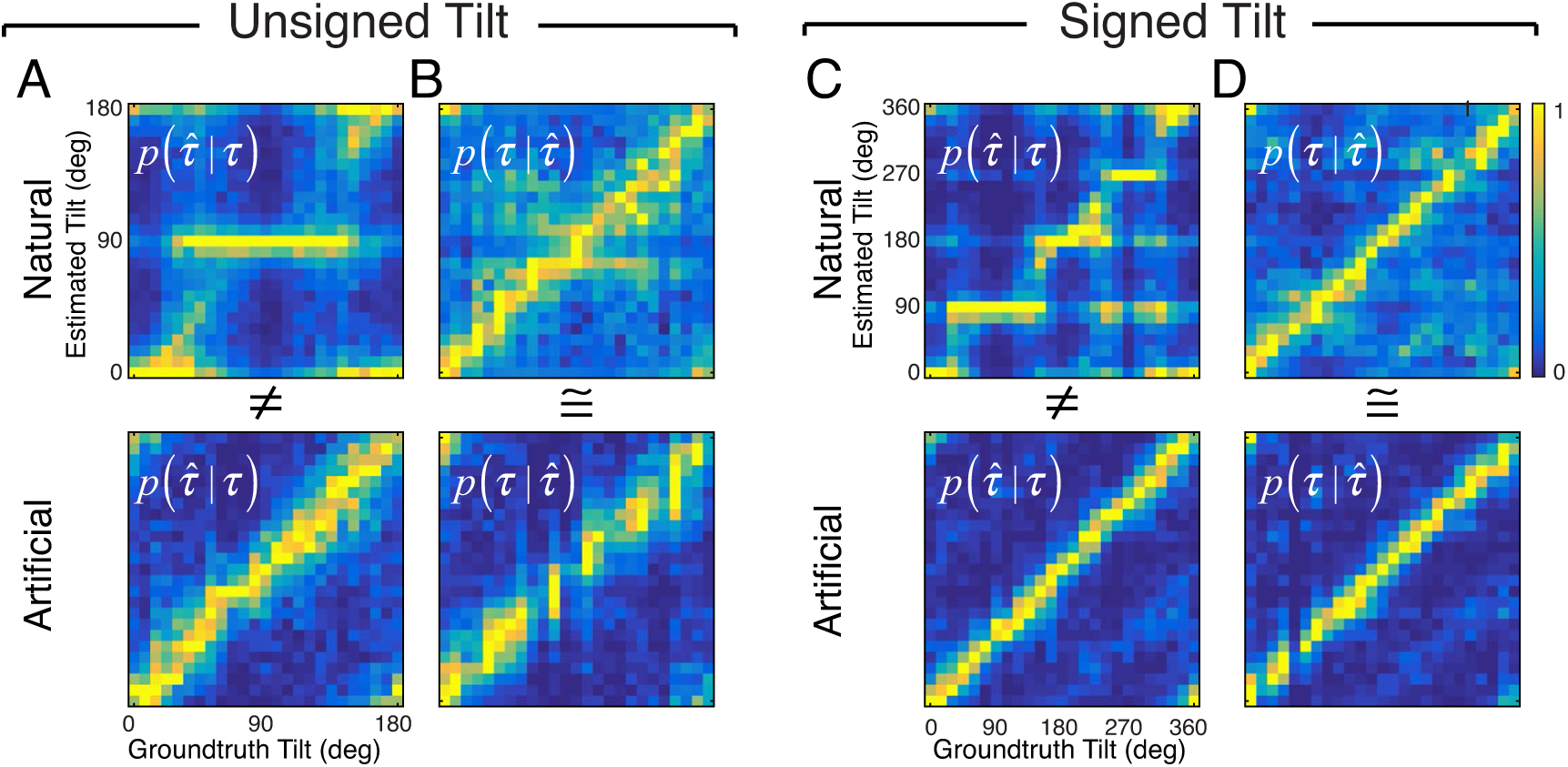
Alternative visualization of data in Figures 3 and 4 in the main text. Top row: Natural stimuli. Bottom row: Artificial stimuli. **A** Conditional probability 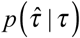 of estimated tilts given groundtruth tilt in the unsigned tit domain: *τ* = [0, 180°) Each column of each subplot represents the conditional probability distribution of estimates for a different groundtruth tilt. Hence, each column represents the data in each subplot of Fig. 3. **B** Conditional probability 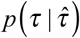 of groundtruth tilts given estimated tilt. Each row of each subplot represents the conditional probability distribution of groundtruth tilts for a different estimate. Hence, each row represents the data in each subplot of Fig. 4. **C,D** Same as A,B but in the signed tilt domain: *τ* = [0°, 360°). For visualization, each conditional probability distribution is normalized so that the maximum probability equals 1.0 (colorbar). The distributions of tilt estimates from natural and artificial stimuli conditional on groundtruth tilts are very different (A,C). In contrast, the distributions of groundtruth tilts conditional on estimates from natural and artificial stimuli are very similar (B,D). Thus, regardless of the stimulus type, the information provided about groundtruth by human tilt estimates is of similar quality. This property of the estimates should simplify subsequent processing that combines local into more global estimates (see main text).

**Figure S5.**
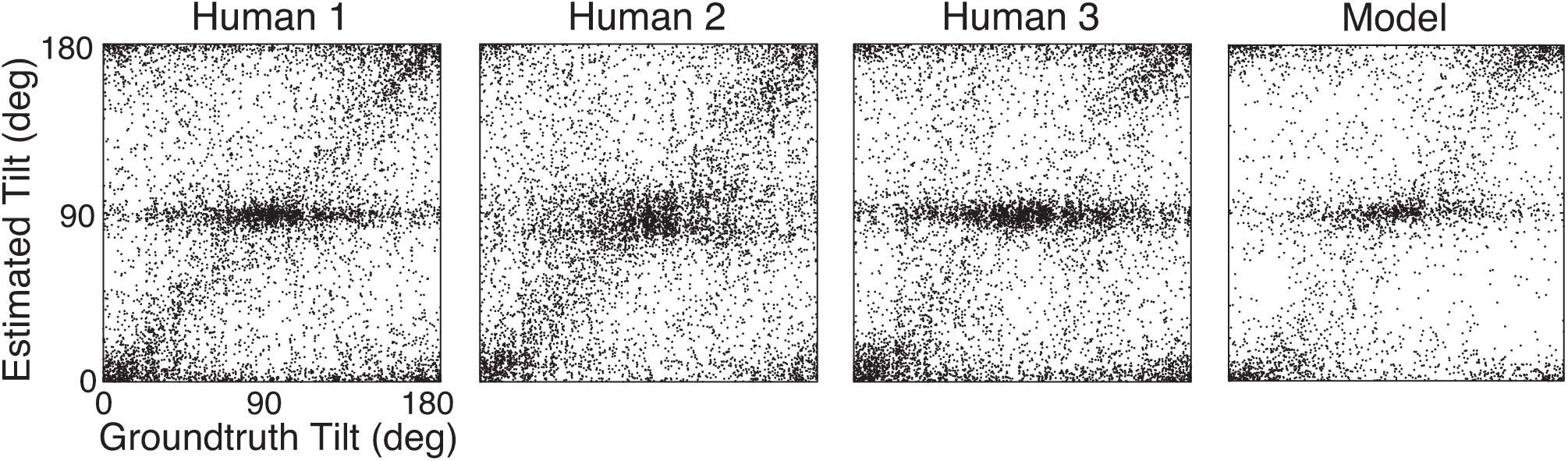
Human and normative model unsigned tilt estimates. Each panel shows the unsigned tilt estimate of a human or model observer plotted against the groundtruth tilt of every stimulus presented in the experiment.

**Figure S6.**
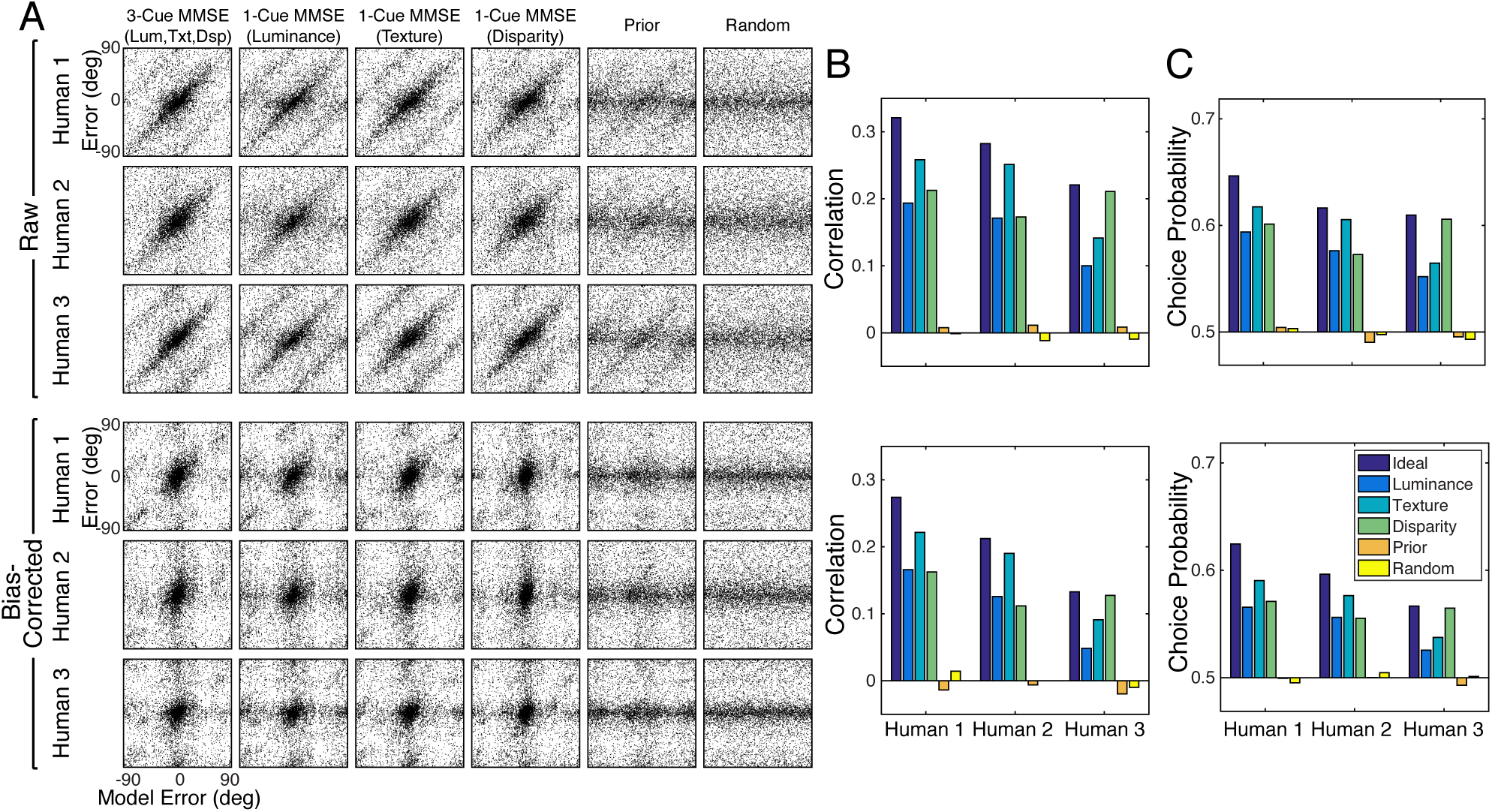
Six alternative models for predicting human tilt estimation performance. Each model sets its estimates equal to (i) the minimum-mean squared error (MMSE) estimates based on three-cues (i.e. the normative model used in the main text); (ii-iv) the MMSE estimates based on each single cue alone (Luminance; Texture; and Disparity); (v) random tilt samples from the tilt prior (Prior); and (vi) random tilt samples from a uniform distribution of tilts (Random) **A** Human trial-by-trial errors plotted against the errors made by each of the models. Upper rows show raw errors; lower rows show bias-corrected errors. **B** Circular correlation coefficient for each of the models considered in A. **C** Choice probability for each of the models considered in A. Here, we define choice probability as the proportion of trials that the sign of the model error predicts the sign of the human errors. The pattern is similar to circular correlation coefficient. The MMSE model based on three image cues predicts humans tilt estimation errors better than all other models. Additionally, we assessed a number of ad hoc models not shown here. Three models that set the tilt estimate equal to each of the single cue values (i.e. cue-gradient orientation) predict human performance more poorly than the three-cue normative model used in the main text, but better the prior or the random model. A model that averages the single-cue values (with equal weights) predicts human estimation better than the single-cue MMSE estimators, but worse than the three-cue normative model used in the main text.

**Figure S7.**
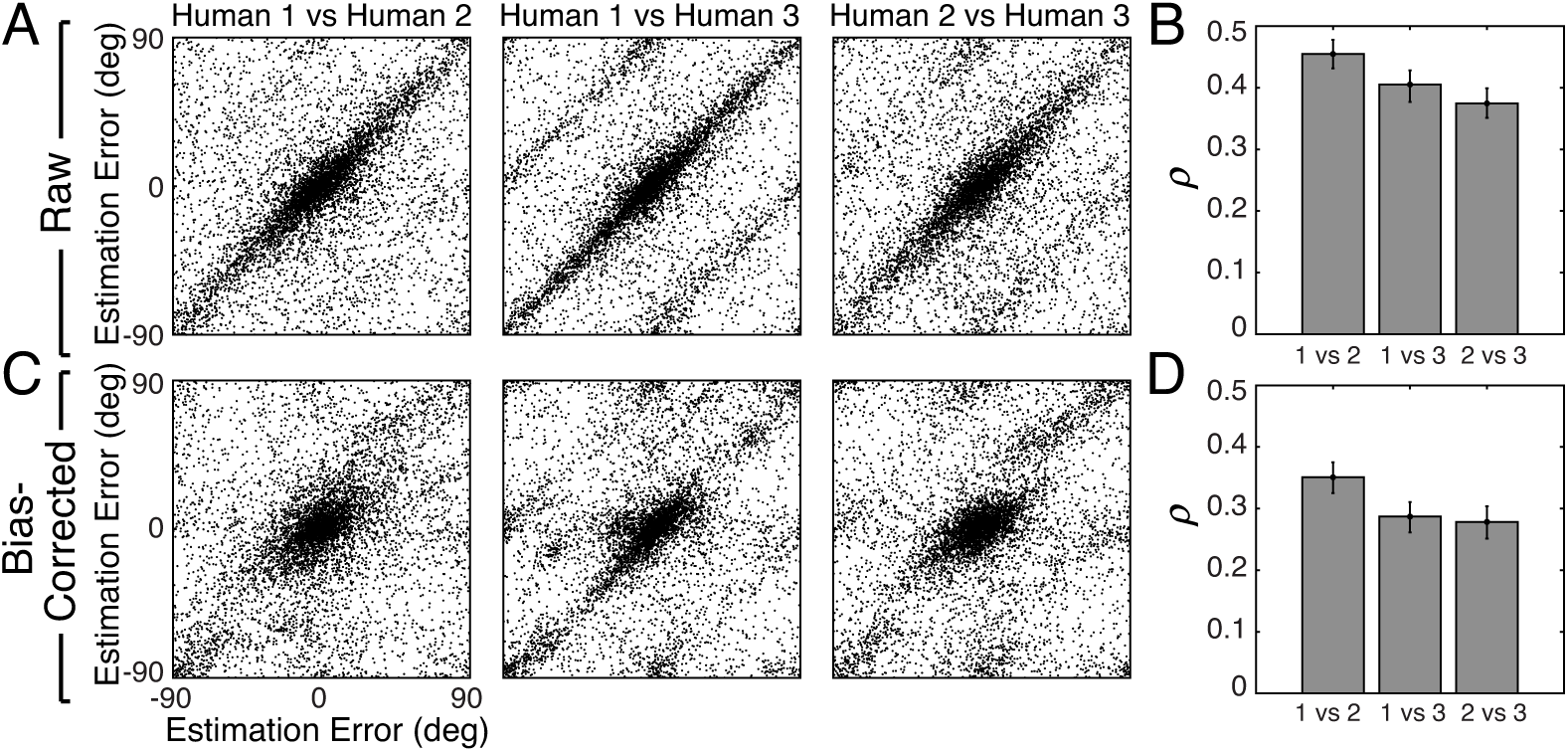
Trial-by-trial estimation errors between humans. The diagonal structure in the plots indicates that trial-by-trial errors are correlated. **A** Raw trial-by-trial errors with natural stimuli between humans. **B** Correlation coefficients (circular) for trial-by-trial errors between humans. The error bars represent 95% confidence intervals from 1000 bootstrapped samples of the correlation coefficient. **C** Bias-corrected errors in natural stimuli. **D** Correlation coefficient for bias-corrected errors.

**Figure S8.**
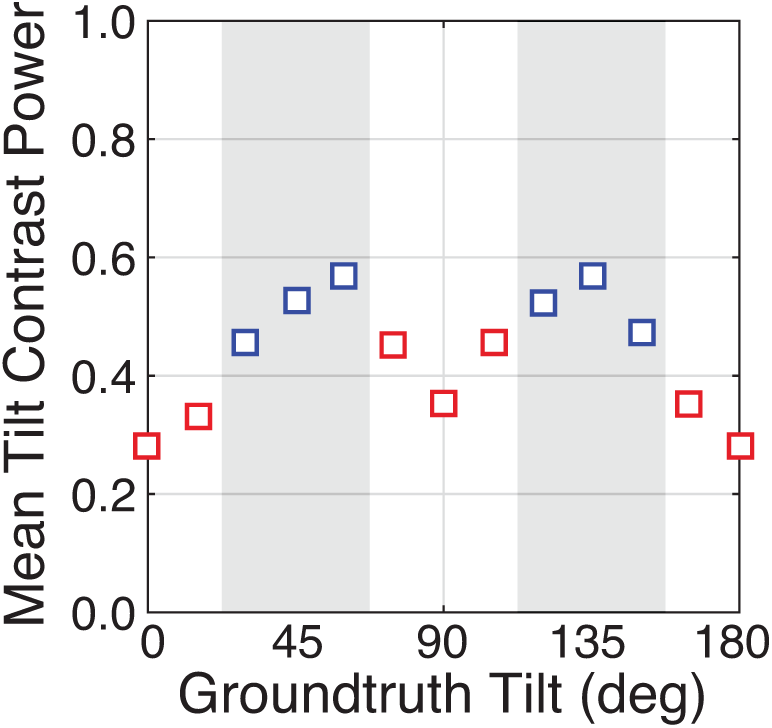
Tilt contrast power co-varies with groundtruth tilt. Oblique groundtruth tilts (e.g. 45° and 135°, shaded areas) have higher tilt contrast powers than cardinal groundtruth tilts (e.g. 0° and 90°). Oblique tilts tend to be associated with less planar (i.e. more bumpy) regions of natural scenes.

**Figure S9.**
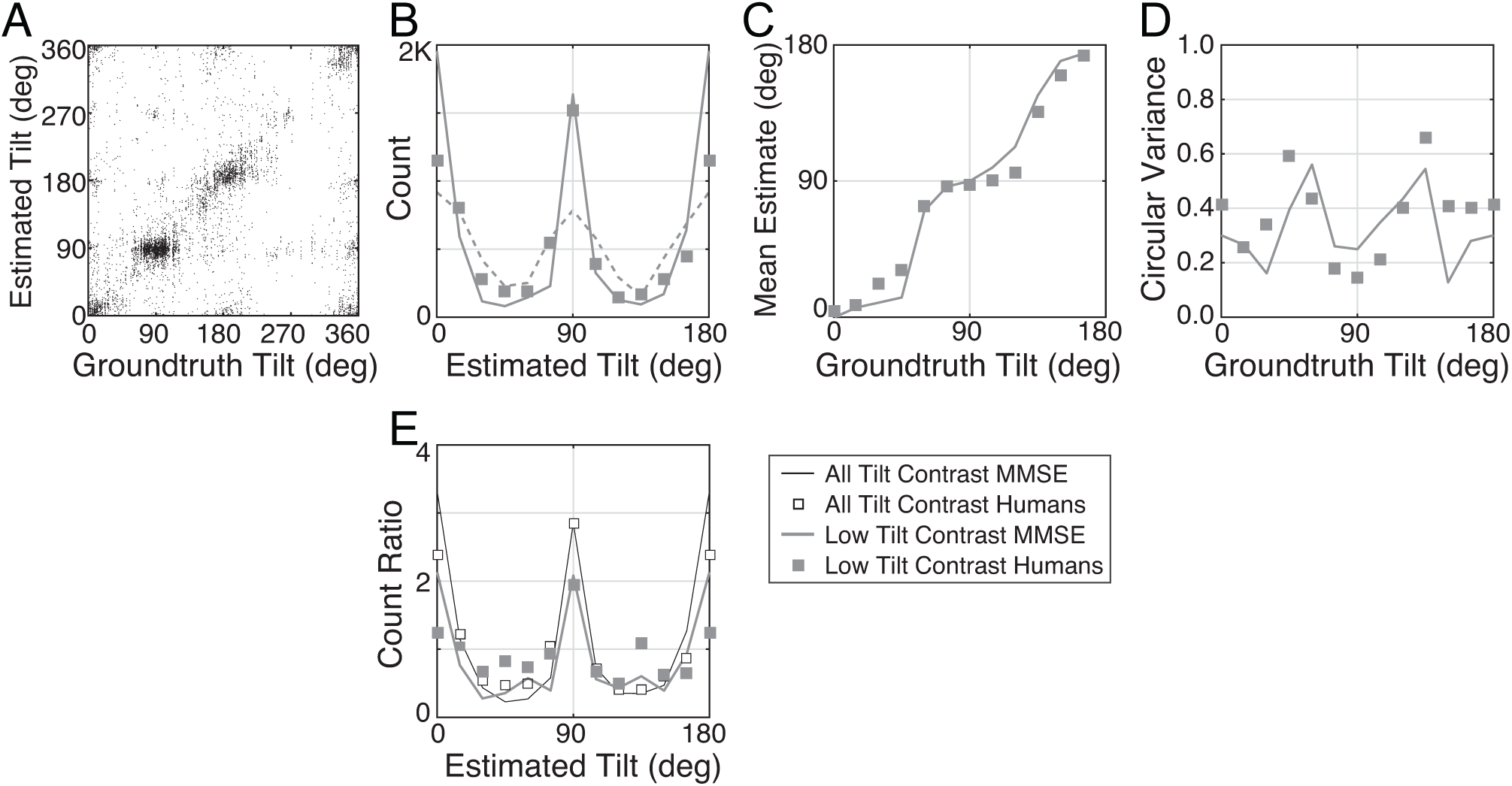
Tilt estimation performance with near-planar natural stimuli. We reanalyzed performance for the subset of natural stimuli with the lowest tilt contrast powers (tilt contrast power ≤ 0.05; bottom quintile). For these near-planar stimuli, the mean tilt estimation error is 16.9°, slightly lower than the mean error with artificial stimuli (18.1°; see Fig. 7A). **A** Raw responses. **B** Histogram of estimates. The dashed horizontal curve shows the distribution of groundtruth tilts in this subset of stimuli, which is not uniform (dashed curve). Unlike with planar artificial stimuli (c.f. Fig. 2E), the histogram of human responses to near-planar natural stimuli over-represents the cardinal tilts relative to the frequency of the presented stimuli. The normative model nicely predicts the histogram of human estimates (solid curve). **C** Mean estimates as a function of groundtruth tilt. **D** Estimate variance as a function of groundtruth tilt. The mean and variance functions are different from the mean and variance functions yielded by planar artificial stimuli. Thus, tilt contrast is not solely responsible for the differences between performance with natural and artificial stimuli. **E** Estimate count ratios (i.e. estimated / presented) at each tilt. With near-planar natural stimuli, cardinal tilts are still estimated much more frequently than with planar artificial stimuli. Tilt contrast is not the only factor responsible for performance differences between natural and commonly used artificial stimuli.

## References

[1] Stevens KA. Slant-tilt: The visual encoding of surface orientation. Biol Cybern 1983;46:183–95. doi:10.1007/BF00336800.

[2] Todd JT, Koenderink JJ, van Doorn AJ, Kappers AM. Effects of changing viewing conditions on the perceived structure of smoothly curved surfaces. J Exp Psychol Hum Percept Perform 1996;22:695–706.

[3] Knill DC. Surface orientation from texture: ideal observers, generic observers and the information content of texture cues. Vision Research 1998;38:1655–82.

[4] Knill DC. Ideal observer perturbation analysis reveals human strategies for inferring surface orientation from texture. Vision Research 1998;38:2635–56.

[5] Hillis JM, Watt SJ, Landy MS, Banks MS. Slant from texture and disparity cues: optimal cue combination. J Vis 2004;4:967–92. doi:10.1167/4.12.1.

[6] Watt SJ, Akeley K, Ernst MO, Banks MS. Focus cues affect perceived depth. J Vis 2005;5:834–62. doi:10.1167/5.10.7.

[7] Burge J, Girshick AR, Banks MS. Visual-haptic adaptation is determined by relative reliability. J Neurosci 2010;30:7714–21. doi:10.1523/JNEUROSCI.6427-09.2010.

[8] Fleming RW, Holtmann-Rice D, Bulthoff HH. Estimation of 3D shape from image orientations. Proc Natl Acad Sci USa 2011;108:20438–43. doi:10.1073/pnas.1114619109/-/DCSupplemental.

[9] Todd JT. The visual perception of 3D shape. Trends Cogn Sci (Regul Ed) 2004;8:115– 21. doi:10.1016/j.tics.2004.01.006.

[10] Marlow PJ, Todorovic D, Anderson BL. Coupled computations of three-dimensional shape and material. Curr Biol 2015;25:R221–2. doi:10.1016/j.cub.2015.01.062.

[11] Rosenholtz R, Malik J. Surface orientation from texture: isotropy or homogeneity (or both)? Vision Research 1997;37:2283–93.

[12] Rosenberg A, Cowan NJ, Angelaki DE. The visual representation of 3D object orientation in parietal cortex. J Neurosci 2013;33:19352–61. doi:10.1523/JNEUROSCI.3174-13.2013.

[13] Murphy AP, Ban H, Welchman AE. Integration of texture and disparity cues to surface slant in dorsal visual cortex. J Neurophysiol 2013;110:190–203. doi:10.1152/jn.01055.2012.

[14] Velisavljevic L, Elder JH. Texture properties affecting the accuracy of surface attitude judgements. Vision Research 2006;46:2166–91. doi:10.1016/j.visres.2006.01.010.

[15] Li A, Zaidi Q. Perception of three-dimensional shape from texture is based on patterns of oriented energy. Vision Research 2000;40:217–42.

[16] Li A, Zaidi Q. Three-dimensional shape from non-homogeneous textures: carved and stretched surfaces. J Vis 2004;4:860–78. doi:10.1167/4.10.3.

[17] Saunders JA, Knill DC. Perception of 3D surface orientation from skew symmetry. Vision Research 2001;41:3163–83.

[18] Welchman AE, Deubelius A, Conrad V, Bülthoff HH, Kourtzi Z. 3D shape perception from combined depth cues in human visual cortex. Nat Neurosci 2005;8:820–7. doi:10.1038/nn1461.

[19] Sanada TM, Nguyenkim JD, DeAngelis GC. Representation of 3-D surface orientation by velocity and disparity gradient cues in area MT. J Neurophysiol 2012;107:2109–22. doi:10.1152/jn.00578.2011.

[20] Tsutsui K, Jiang M, Yara K, Sakata H, Taira M. Integration of perspective and disparity cues in surface-orientation-selective neurons of area CIP. J Neurophysiol 2001;86:2856–67.

[21] Burge J, McCann BC, Geisler WS. Estimating 3D tilt from local image cues in natural scenes. J Vis 2016;16:2. doi:10.1167/16.13.2.

[22] Yang Z, Purves D. A statistical explanation of visual space. Nat Neurosci 2003;6:632– 40.

[23] Adams WJ, Elder JH, Graf EW, Leyland J, Lugtigheid AJ, Muryy A. The Southampton-York Natural Scenes (SYNS) dataset: Statistics of surface attitude. Nature Publishing Group 2016;6:35805. doi:10.1038/srep35805.

[24] McDermott J. Psychophysics with junctions in real images. Perception 2004;33:1101– 27. doi:10.1068/p5265.

[25] Knill DC, Richards W. Perception as Bayesian Inference. New York: Cambridge University Press; 1996.

[26] Appelle S. Perception and discrimination as a function of stimulus orientation: The “oblique effect” in man and animals. Psychological Bulletin 1972;78:266. doi:10.1037/h0033117.

[27] Furmanski CS, Engel SA. An oblique effect in human primary visual cortex. Nat Neurosci 2000;3:535–6. doi:10.1038/75702.

[28] Girshick AR, Landy MS, Simoncelli EP. Cardinal rules: visual orientation perception reflects knowledge of environmental statistics. Nat Neurosci 2011;14:926–32. doi:10.1038/nn.2831.

[29] Ernst MO, Banks MS. Humans integrate visual and haptic information in a statistically optimal fashion. Nature 2002;415:429–33. doi:10.1038/415429a.

[30] Burgess AE, Wagner RF, Jennings RJ, Barlow HB. Efficiency of human visual signal discrimination. Science 1981;214:93–4. doi:10.1080/00324728.2016.1201588.

[31] Yang Z, Purves D. Image/source statistics of surfaces in natural scenes. Network 2003;14:371–90.

[32] Land MF, Hayhoe M. In what ways do eye movements contribute to everyday activities? Vision Research 2001;41:3559–65.

[33] Pelz JB, Rothkopf C. Oculomotor behavior in natural and man-made environments. In: van Gompel RPG, Fischer M, Murray WS, Hill RL, editors. Eye Movements: A Window on Mind and Brain, Eye Movements: A window on Mind and Brain; 2007.

[34] Dorr M, Martinetz T, Gegenfurtner KR, Barth E. Variability of eye movements when viewing dynamic natural scenes. J Vis 2010;10:28. doi:10.1167/10.10.28.

[35] Geisler WS, Najemnik J, Ing AD. Optimal stimulus encoders for natural tasks. J Vis 2009;9:17.1–16. doi:10.1167/9.13.17.

[36] Burge J, Jaini P. Accuracy Maximization Analysis for Sensory-Perceptual Tasks: Computational Improvements, Filter Robustness, and Coding Advantages for Scaled Additive Noise. PLoS Comput Biol 2017;13:e1005281. doi:10.1371/journal.pcbi.1005281.

[37] Burge J, Geisler WS. Optimal defocus estimation in individual natural images. Proc Natl Acad Sci USa 2011;108:16849–54. doi:10.1073/pnas.1108491108.

[38] Burge J, Geisler WS. Optimal defocus estimates from individual images for autofocusing a digital camera, Proceedings of SPIE; 2012. doi:10.1117/12.912066.

[39] Burge J, Geisler WS. Optimal disparity estimation in natural stereo images. J Vis 2014;14. doi:10.1167/14.2.1.

[40] Burge J, Geisler WS. Optimal speed estimation in natural image movies predicts human performance. Nat Commun 2015;6:7900. doi:10.1038/ncomms8900.

[41] Marr D. Vision. New York: W H Freeman & Company; 1982.

[42] Oruç I, Maloney LT, Landy MS. Weighted linear cue combination with possibly correlated error. Vision Research 2003. doi:10.1016/S0042-6989(03)00435-8.

[43] Burge J, Fowlkes CC, Banks MS. Natural-scene statistics predict how the figure-ground cue of convexity affects human depth perception. J Neurosci 2010;30:7269– 80. doi:10.1523/JNEUROSCI.5551-09.2010.

[44] Weiss Y, Simoncelli EP, Adelson EH. Motion illusions as optimal percepts. Nat Neurosci 2002;5:598–604. doi:10.1038/nn858..

[45] ocker AA, Simoncelli EP. Noise characteristics and prior expectations in human visual speed perception. Nat Neurosci 2006;9:578–85. doi:10.1038/nn1669.

[46] Tyler CW, Julesz B. Binocular cross-correlation in time and space. Vision Research 1978;18:101–5.

[47] Banks MS, Gepshtein S, Landy MS. Why is spatial stereoresolution so low? J Neurosci 2004;24:2077–89. doi:10.1523/JNEUROSCI.3852-02.2004.

[48] Clerc M, Mallat S. The texture gradient equation for recovering shape from texture. Pattern Analysis and Machine Intelligence 2002.

[49] Galasso F, Lasenby J. Shape from Texture: Fast Estimation of Planar Surface Orientation via Fourier Analysis. Bmvc 2007.

[50] Malik J, Rosenholtz R. Computing local surface orientation and shape from texture for curved surfaces. Int J Comput Vis 1997.

[51] Massot C, Hérault J. Model of frequency analysis in the visual cortex and the shape from texture problem. Int J Comput Vis 2008. doi:10.1007/s11263-007-0048-x.

